# Impact of Sexual Transmission to Sex-specific Attack Rates in Zika Epidemics

**DOI:** 10.1101/347104

**Authors:** Ana Carolina W.G. de Barros, Kaline G. Santos, Eduardo Massad, Flávio Codeço Coelho

## Abstract

In 2015 and 2016 South America went through the largest Zika epidemic in recorded history. One important aspect of this epidemic was the impact on newborns due to the effect of Zika on development of the central nervous system leading to severe malformations. Another aspect of the Zika epidemic which became evident from the data was the importance of the sexual route of transmission leading to increased risk for women. Here propose a mathematical model for the transmission of the Zika virus including sexual transmission via all forms of sexual contact, as well as simplified vector transmission, assuming a constant availability of mosquitoes. From this model we derive an expression for *ℛ*_0_ which can be used to study and analyze the relative contributions of the different routes of Zika transmission and the male to female sexual transmission route vis-a-vis vectorial transmission. We also fit the model to data from the 2016 Zika epidemic in Rio de Janeiro, to estimate the values of key parameters of the model.

## Introduction

The Zika virus (ZIKV) originated from the Zika forest in Uganda where was discovered for the first time in a rhesus monkey in 1947^1^. The first cases of human infection were recorded in Nigeria and Tanzania from 1952 to 1954^2^, spreading slowly across the Asian continent. Before 2007, ZIKV was not considered a disease of substantial concern to human beings because only isolated cases involving small populations had been reported worldwide^3^. The ZIKV transmission that had previously only been documented in regions of Africa and Asia, in 2007 it was detected in Yap, Micronesia, causing a small outbreak. Forty-nine ZIKV infected cases were confirmed. In 2012/2013, it caused a new outbreak in French Polynesia and spreading across the others Pacific islands, resulting in an epidemic with more than 400 confirmed cases^4^.

In March of 2015, the virus was detected for the first time in Brazil^5^. In October 2015, a growing number of cases of newborns with microcephaly were reported in Pernambuco. In November, after confirmation of the ZIKV presences in the amniotic fluid of pregnant women in the state of Paráiba, the association between the virus infection and microcephaly was confirmed^2^.

ZIKV spread rapidly throughout Brazil and in less than a year it had reached the entire country, spreading to neighboring countries. It has been estimated that in 2015 alone there were between 440,000 and 1,300,000 Zika cases in Brazil, resulting in the largest ZIKV epidemic reported so far^6^.

Although ZIKV is a flavivirus transmitted to humans primarily through the bite of infected Aedes mosquitoes^7^ it can also spread through sexual contact. Studies have shown the presence of a infectious viral load in the semen after Zika infection^8^ indicating that we are dealing with a pathogen that is also sexually transmissible^9^. Although not very common, other forms of transmission between humans where confirmed recently from the identification of virus in the urine and saliva^10^. The most reported and well characterized form of sexual transmission among humans if from male to female. However, transmission through homosexual relations and from women to men have also been reported^11,12^.

Zika transmission has been modeled mathematically before^13–15^. To our best knowledge, none of the published models explicitly include all the sexual transmission modes and derives an analytical expression for RO reflecting them.

Given the current understanding of mechanisms behind the transmission of ZIKV and its consequences to human health we expect our model to help in the investigation of the relative importance of the vectorial and sexual transmission to the dynamics of Zika.

Finally, we fit the model to data from the 2016 Zika epidemic in Rio de Janeiro to estimate some of its parameters, and to demonstrate that this model is able to explain a real Zika epidemic with the different burdens for women and men.

## Methods

The proposed epidemic model used is an adaptation of a SEIR model proposed earlier^16^. We assume a constant population size, ignoring demographic processes such as birth, death and immigration as well as seasonal fluctuations.

At any given instant *t*, the state of the system is represented by the fractions of the total population *N*(*t*) in each of the immunological states described in equations 1, further split by sex. The dynamics of the mosquito population is not explicitly included in the model being represented by a single parameter of vectorial transmission.

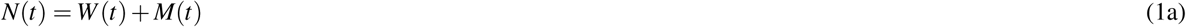

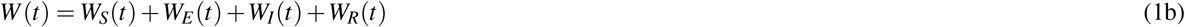

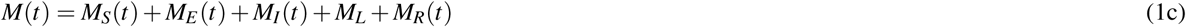

The susceptible female and male populations, *W*_*S*_(*t*) and *M*_*S*_(*t*), respectively are exposed to the Zika virus with a *β*_*S*_ sexual transmission rate and a *β*_*V*_ vector transmission rate. Since we are not modeling the mosquito population dynamics we simplify vectorial transmission to a mass-action term, e.g. *β*_*V*_ (*M*_*I*_ (*t*) + *W*_*I*_ (*t*))*W*_*S*_(*t*).

Due to the fact that the virus remain viable in the semen, men may go into a latent state, *M*_*L*_(*t*) indicating a longer sexual infectious period as a whole. However, since the latent period is not the same for all men we add the parameter *ρ* that takes into consideration the fact that not all men go into latency.

To account for different rates of sexual transmission between sexes, we take the male-to-female transmission rate to be the baseline, the most effective, due to the larger inoculum. We denote it by *β*_*S*_ and define *k*_*WW*_, *k*_*WM*_, *k*_*MM*_, and *k*_*L*_, to be a ratio of this baseline transmission contributed by other modes of sexual transmission, respectively woman-to-woman, woman-to-man, man-to-man and latent-man-to-woman. We further assume these ratios to range from 0 to 1, meaning the other modes can at most be as effective as the male-to-female transmission.

We assume permanent and perfect immunity against new ZIKV infections after the first infection, therefore, recovered individual are not included explicitly in the model.

The proposed model consists of a system of seven ordinary differential equations:

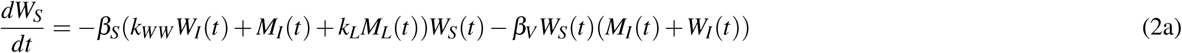

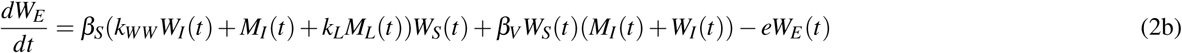

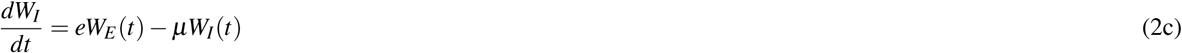

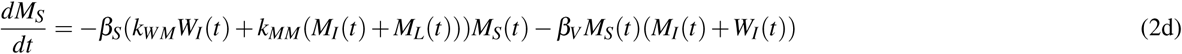

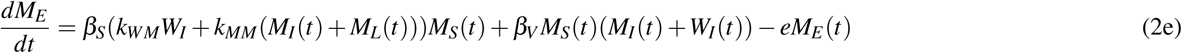

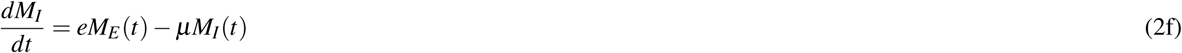

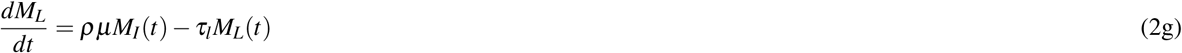

The model’s variables and parameters are described in table 1.

**Table 1.** Variables and parameters of the model. Values obtained from the literature are marked with references numbers. Ranges marked with a ^†^ correspond to values explored in simulations, but for which no experimental data could be found.^*\**^ fraction of the entire population, *N*

| Symbols | Description | Value Range |
| --- | --- | --- |
| $W_S$ | Susceptible women | $[0, 1]^*$ |
| $W_E$ | Exposed women | $[0, 1]^*$ |
| $W_I$ | Infectious women | $[0, 1]^*$ |
| $M_S$ | Susceptible men | $[0, 1]^*$ |
| $M_E$ | Exposed men | $[0, 1]^*$ |
| $M_I$ | Infectious men | $[0, 1]^*$ |
| $M_L$ | Latent men | $[0, 1]^*$ |
| $\mu$ | recovery rate $[\text{days}^{-1}]$ | $[.001, .1]^{14}$ |
| $\beta_V$ | vector transmission rate $[\text{days}^{-1}]$ | $[.1, .75]^{14}$ |
| $\beta_S$ | sexual transmission rate $[(\text{people} \times \text{days})^{-1}]$ | $[0, 2]^{\dagger}$ |
| $k_{WW}$ | women to women transmissibility modifier | $[0, 1]^{\dagger}$ |
| $k_{WM}$ | women to men transmissibility modifier | $[0, 1]^{\dagger}$ |
| $k_{MM}$ | men to men transmissibility modifier | $[0, 1]^{\dagger}$ |
| $k_L$ | latent period transmissibility modifier | $[0, 1]^{\dagger}$ |
| $e$ | incubation rate $[\text{days}^{-1}]$ | $[0.14, .5]^{14}$ |
| $\rho$ | fraction of men becoming latent | $[0, 1]^{\dagger}$ |
| $\tau_l$ | latent recovery rate $[\text{days}^{-1}]$ | $[0.025, 0.1]^8$ |

### Attack Rates by Sex

From the model defined by equations 2 we can define another two differential equations to track the Attack rates over time. In epidemiology, the attack rate, is the ratio between the number of cases and the population at risk. It is usually calculated for the entire epidemic, but we can also calculate it up to each point in time since the beginning of the epidemic, from the solutions of the following equations for the accumulated number of cases.

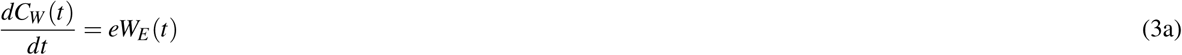

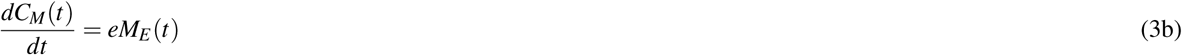

Since we are working with normalized populations where *N*(*t*) = 1, ∀*t* with equal number of men and women, the attack ratios for women and men are given by *AR*_*W*_ (*t*) = 2*C*_*W*_ (*t*) and *AR*_*M*_(*t*) = 2*C*_*M*_(*t*), respectively. The ratio *AR*_*W*_ (*t*)*/AR*_*M*_(*t*), will represent the female burden relative to men. To calculate this ratio which is shown on figure 5, we have fixed an epidemic duration of 120 days to accumulate cases. We chose 120 days because arbovirus epidemics are usually restricted by weather conditions affecting mosquito activity and longevity and rarely last longer than 120 days.

### Sexual Force of infection

As we are studying the impact of sexual transmission to transmission of Zika, it is worth taking a closer look at the sexual force of infection. The force of infection, in epidemic models is typically defined as the product of the transmissibility parameter and the fraction of infectious individuals in the populations. Here since infection can happen both through direct sexual contact and vector transmission, it makes sense to distinguish between a sexual force of infection and a vectorial one. Let’s first define the sexual force of infection for each sex, since their exposure to sexual infection is different. Let *λ*_*SW*_ denote the force of infection of women and *λ*_*SM*_ the force of infection of men.

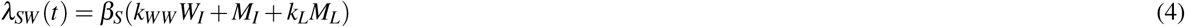

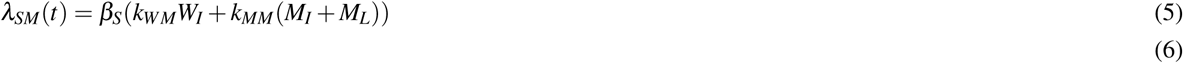

If we assume that transmission from men to women is the only relevant form of sexual transmission(*k*_*WW*_ = *k*_*WM*_ = *k*_*MM*_ = 0), we can simplify the sexual forces of infections above to *λ*_*SW*_ = *β*_*S*_(*M*_*I*_ + *K*_*L*_*M*_*L*_) and *λ*_*SM*_ = 0.

We can also define a vectorial force of infection in a similar fashion: *λ*_*V*_ = *β*_*V*_ (*M*_*I*_ + *W*_*I*_). Then we can rewrite our model as

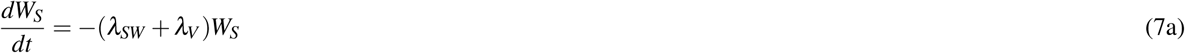

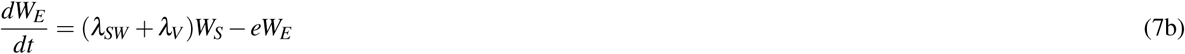

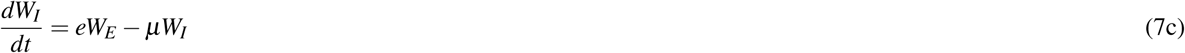

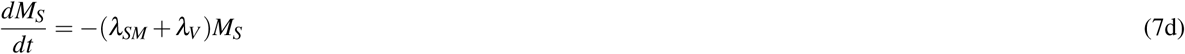

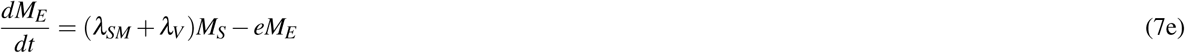

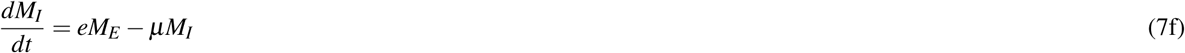

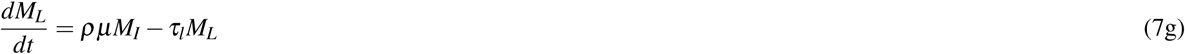

### Fitting the model to data

In order to approximate the relative relative contributions of sexual and vectorial transmission in the context of the proposed model. We fitted to model to the observed prevalence of Zika in men and women in the 2016 epidemic in Rio de Janeiro^10^. We calculated the prevalence of Zika for both sexes by dividing the weekly reported number of cases by the population size for each sex multiplied by the underreporting rate estimated for that epidemic^17^.

To fit the model to data we applied the Bayesian inference methodology proposed by Coelho et al.^18^.

## Results

We performed numerical simulations of the dynamics based on parameter values obtained from the literature. For the parameters were no experimental measurements where available, we explore ranges of values described on table 1.

Figure 1 shows one of the simulations where it is interesting to notice the higher prevalence of both exposed and infectious women in the first 2-3 months of the epidemic.

**Figure 1.**
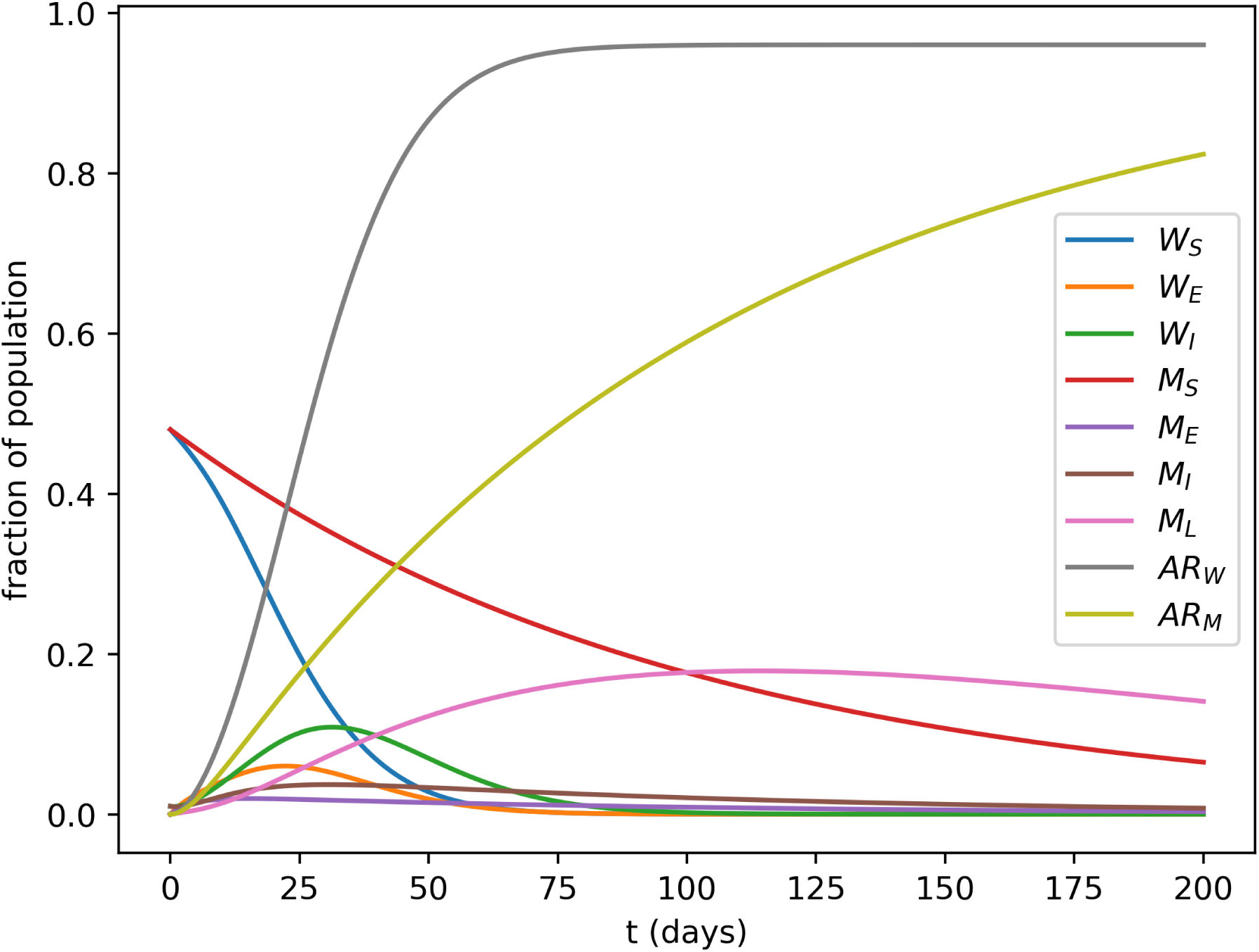
Simulation of the model’s dynamics with *β*_*s*_ = 0.25, *β*_*v*_ = 0.01, *µ* = 0.1, *e* = 0.2, *τ*_*l*_ = 0.01, *ρ* = 1 and *K*_*L*_ = 1. *ℛ*_0_ = 3.04 which compatible with values reported by Villela et al.^21^. *AR*_*W*_ and *AR*_*W*_, curves correspond to the attack rates over time for women and men, respectively.

### Sexual force of infection

Due to its dependence on the prevalence of infectious men in the population, the sexual force of infection in women displays a quite different profile as shown in figure 2 in comparison to the vectorial force of infection.

**Figure 2.**
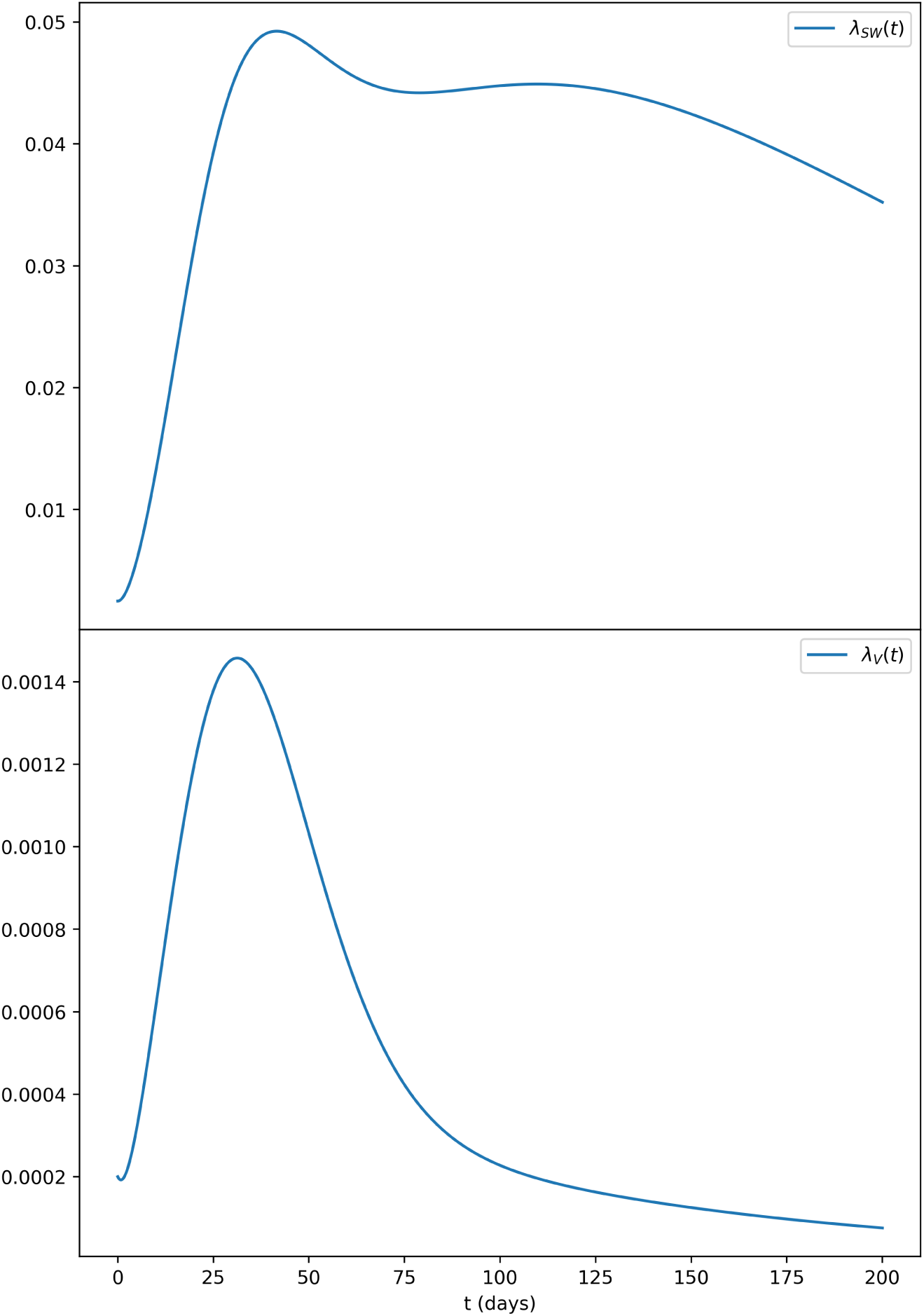
Sexual Force of infection for women with the same parameters as those of figure 1. In the top panel we can observe that the sexual force of infection of women (*λ*_*SW*_ (*t*)) remais elevated for quite a longer period of time if compared to the vectorial force of infection (*λ*_*V*_ (*t*)) shown on the lower panel.

The effects of the sexual force of infection in terms of how effective is the sexual transmission is in the post-viremic stage (*K*_*L*_) can be seen on figure 3.

**Figure 3.**
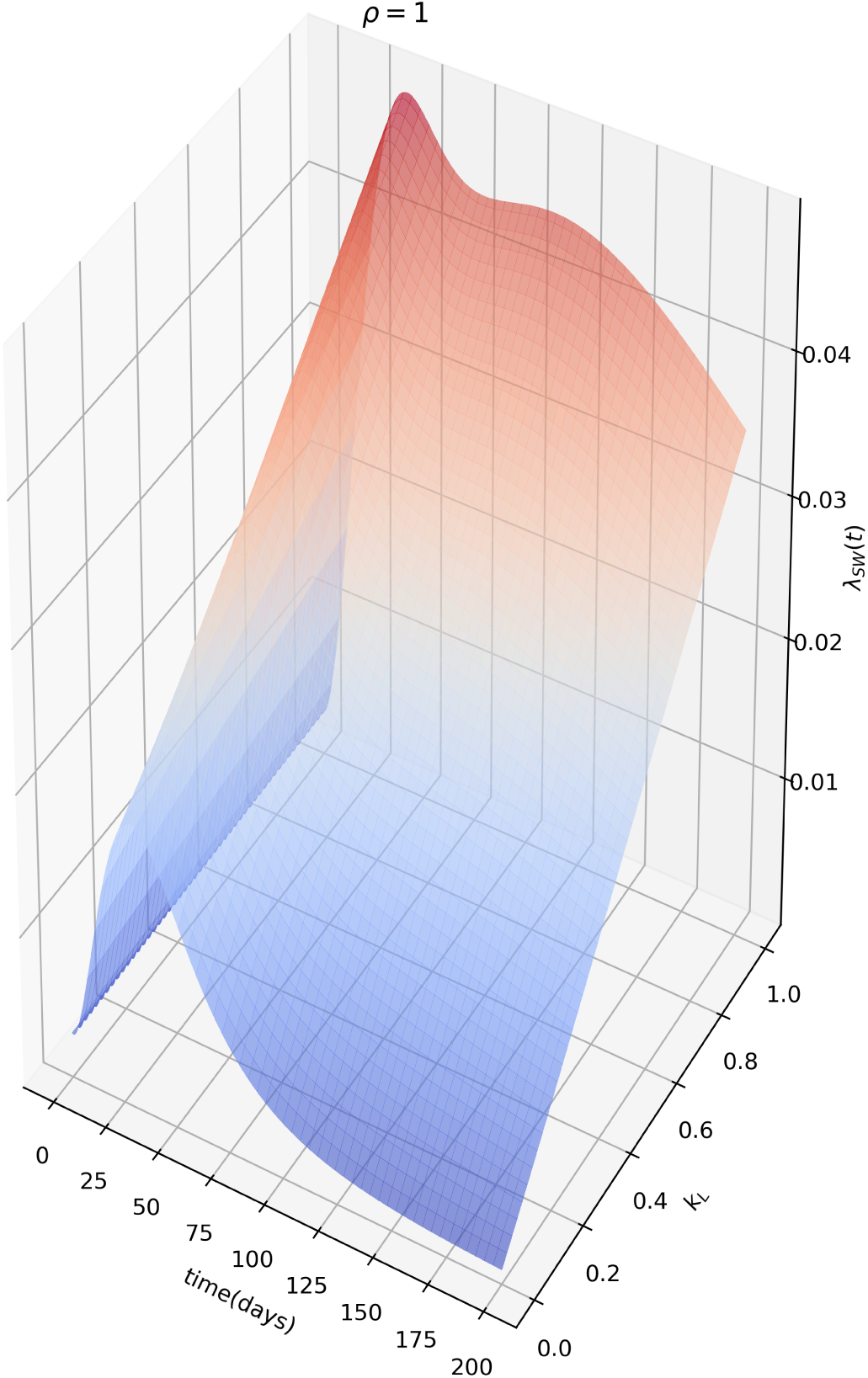
Sexual force of infection as a function of *k*_*L*_, or how effective is the sexual transmission from men to women in the post-viremic phase. Notice that in the absence of effective longer-term sexual transmission from men to women, the dynamics reverts to that of a standard vector borne infection.

Figure 4, depicts the qualitative difference in prevalence dynamics between the effects of underreporting and sexual transmission, the underreported curve for men is obtained by applying a constant 50% reporting rate to the men’s prevalence curve, *M*_*I*_.

**Figure 4.**
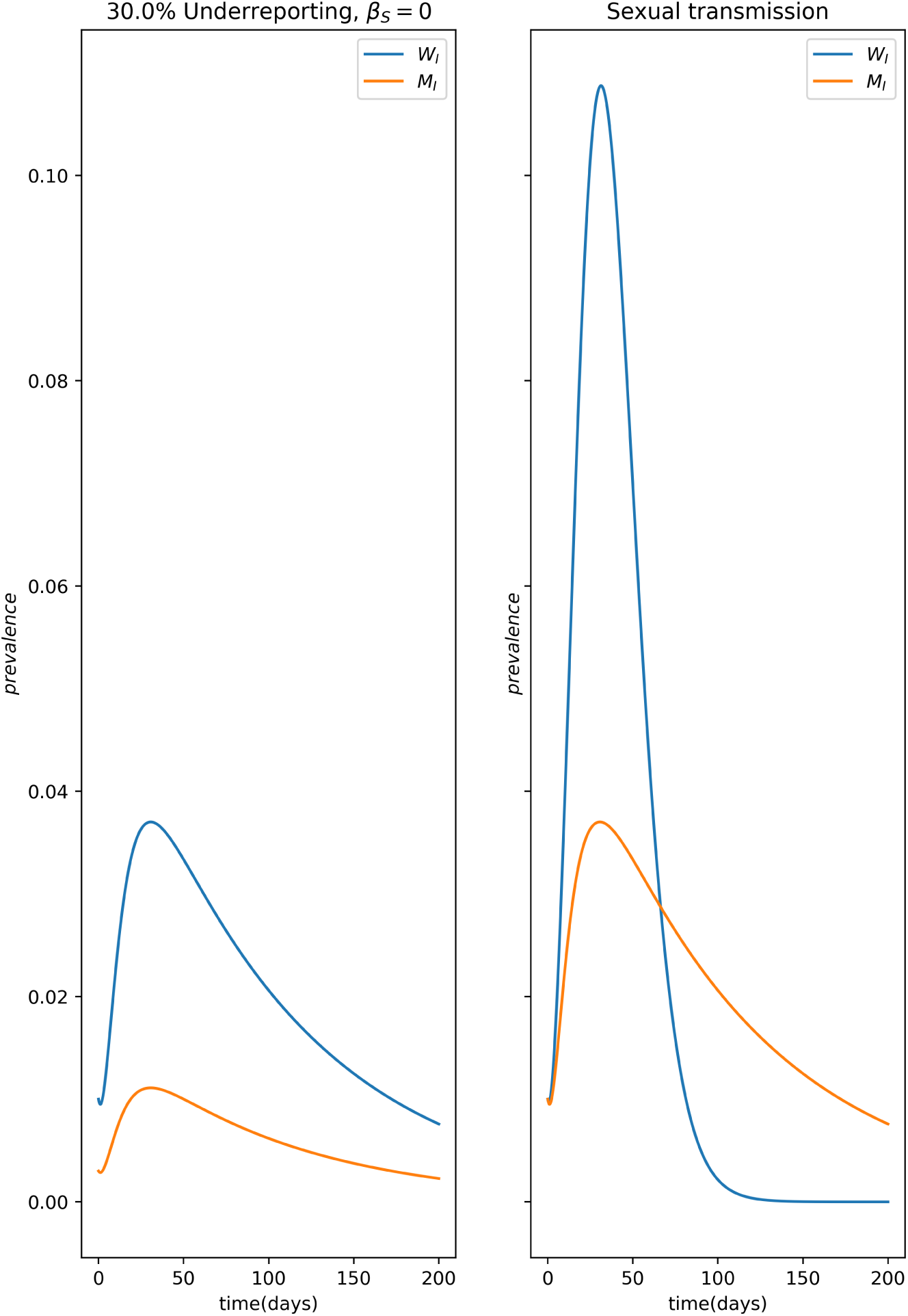
Qualitative differences between the impact of sexual bias in reporting, namely underreporting of male cases (left panel) and sexual transmission in the prevalence curves *W*_*I*_ (*t*) and *M*_*I*_ (*t*) (right panel). Notice that the crossing of the prevalence curves indicates the presence of sexual transmission as this can never happen from underreporting alone.

### The basic reproduction number: *ℛ*_0_

A very important parameter in epidemiology is the basic reproduction number or *ℛ*_0_ of the disease which can be derived from the transmission model. It determines the epidemic potential of a transmissible disease. It is of the average number of infections an infected individual is capable of produce when introduced in a completely susceptible population.

We derive the basic reproduction number for our model by means of the next generation matrix method^19^. According to this method, first we need to distinguish new infections from all the other changes in population. Then we let *m* denote the number of compartments containing infected individuals. They are *W*_*E*_, *W*_*I*_, *M*_*E*_, *M*_*I*_ and *M*_*L*_, so *m* = 5. For clarity we will order the *n* = 7 compartments like so: [*W*_*E*_, *W*_*I*_, *M*_*E*_, *M*_*I*_, *M*_*L*_,*W*_*S*_, *M*_*S*_], separating the *m* = 5 first compartments from the rest. Then we define the vector *ℱ* [*i*] as the rates of appearance of new infections of new infections at each compartments *i, i* = 1, *…, m*, when the system is at a disease-free state. It is worth pointing out that the transfers between exposed to infected and latent (in men cases) are not considered new infections but the progression of an infected individual through many compartments. Likewise we define the vector *𝒱* [*i*] as the net flow of individuals in and out the *m* compartments by other means. Therefore,

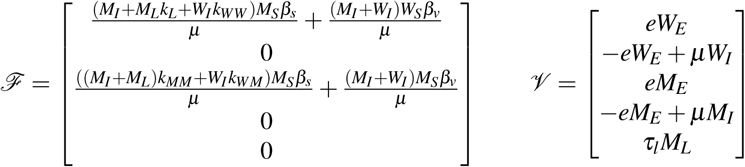

The next step is to calculate the Jacobian of each matrix above, to obtain *F* and *V* at the disease-free equilibrium solution: *W*_*I*_ = 0, *M*_*I*_ = 0, *W*_*S*_ = 1*/*2, *M*_*S*_ = 1*/*2, *W*_*E*_ = 0, *M*_*E*_ = 0 and *M*_*L*_ = 0.

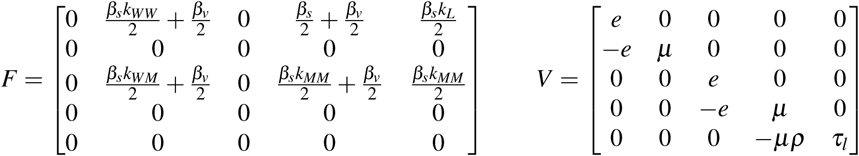

The next generation matrix for the model is given by the product *FV*^-1^ and the *ℛ*_0_ is the spectral radius of resulting matrix.

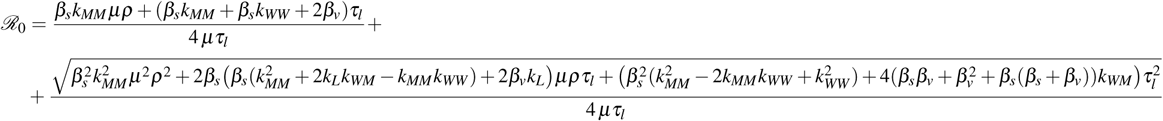

If we disregard Zika sexual transmission between individuals of the same sex and from women to men, we get a simpler expression for *ℛ*_0_

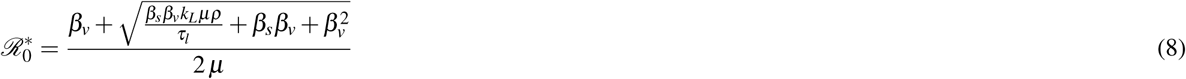

We can also look to the basic reproduction number without sexual transmission, *β*_*s*_ = 0

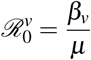

Due to the natural deviation of sexual transmission from a basic mass-action contact rate, as stated in the model, we should apply a correction to 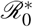 to accommodate for the distribution of the number of sexual partners men has (since we are only considering as sexually infectious). According to Anderson and May^20^(chapter 11, page 233), this correction factor *c* is given by the expression *c* = *m* + *s*^2^*/m*, where *m* is the mean number of sexual partners in the population, and *s*^2^ is its variance. We found the value of *c* = 2.154 using male partner distribution data (*m* = 1.339, *s*^2^ = 1.0917) from a sexual behaviour survey (Carmita Abdo, personal communication). All simulations where done with this correction.

Based on the reduced 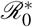 derived from the model, we investigated numerically the importance of the sexual transmission depending on attack ratio for men, *AR*_*M*_, and women, *AR*_*W*_, given different magnitudes of *β*_*s*_ and *β*_*v*_. For these simulations, we disregarded homosexual transmission and from women to men, assuming their contribution are minimal. On figure 5, we can see the ratio 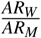 as a function of the two modes of transmission. The green line represents 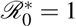 calculated from equation 8. We can see that for more intense values of sexual transmission and moderate vector transmission levels, the total number of female cases is much larger than male cases.

**Figure 5.**
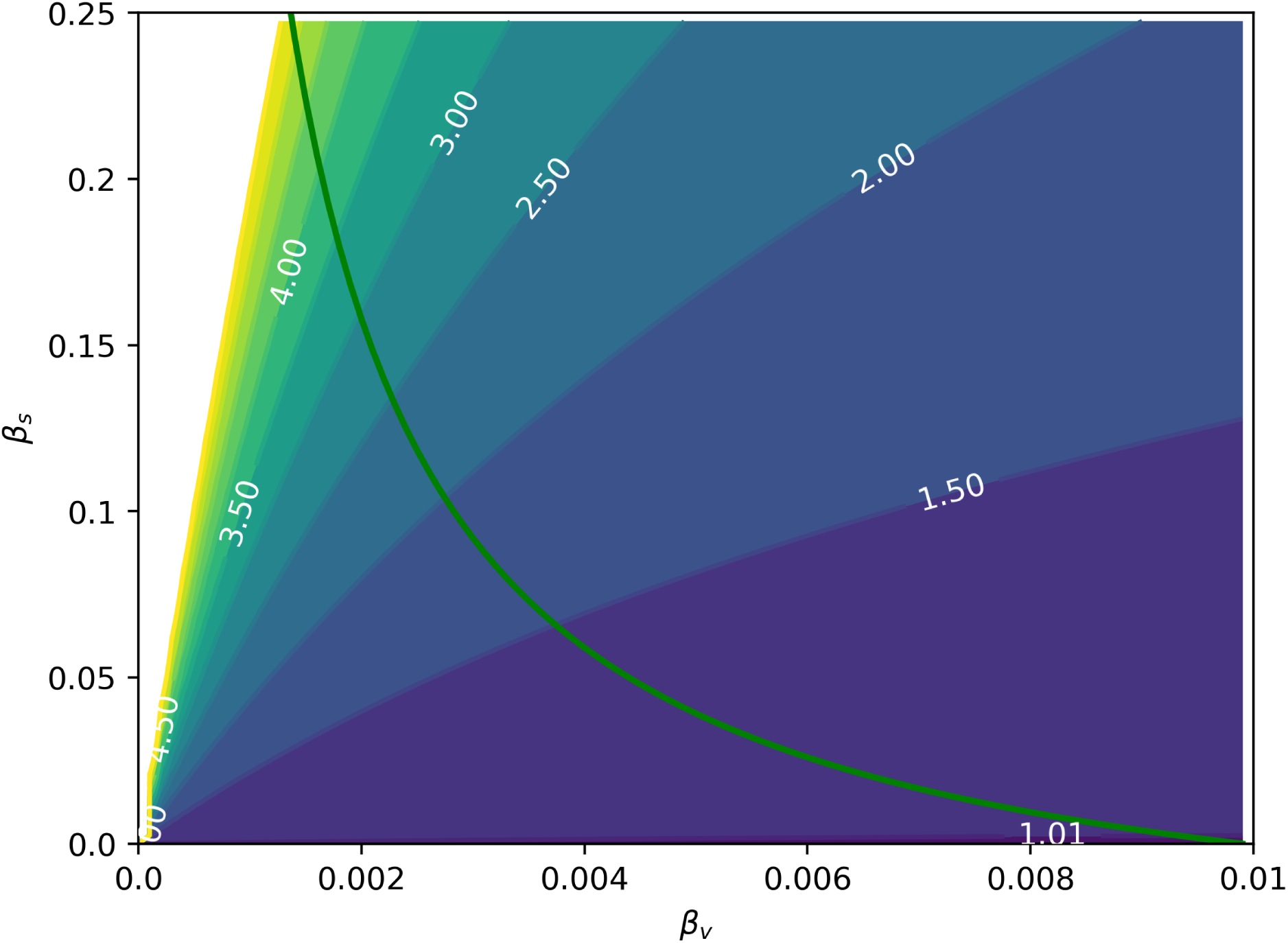
Ratio 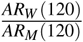 for a range of *β*_*s*_ and *β*_*v*_ values. The green line represents *ℛ*_0_ = 1, i.e. the epidemic threshold. Any point to the right of this curve, has *ℛ*_0_ > 1. It is worth noticing that the reported excess cases reported for Zika in women are possible both during epidemics and off-season.

### Fitting the model to data

Table 2 contains the details about the parameters estimated. The remaining parameters where kept constant at theses values: *k*_*L*_ = 1, *e* = 3.18, *ρ* = 1 and *k*_*MM*_ = *k*_*MH*_ = *k*_*HH*_ = 0. On figure 7, we can see the joint posterior distribution of *β*_*s*_ and *β*_*v*_. On figure 8 we can find the posterior boxplot for all parameters estimated. For the inference, a new parameter, *ur*, was added corresponding to the underreporting of Zika in men relative to women: 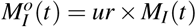 with 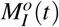 denoting the observed number of male cases. The posterior distributions of the prevalence series is shown on figure 9.

**Table 2.** Parameters estimated from data, along with prior and posterior distribution specifications.^†^ The exponential distribution is parameterized in standardized form (loc, scale). Posterios are given as medians and 95% credible interval. All remaining parameters were kept constant.

| Parameter | Prior | Posterior |
| --- | --- | --- |
| $\beta_s$ | $\mathcal{N}(0.98, 0.3)$ | 0.99[0.02, 1.54] |
| $\beta_v$ | $\mathcal{N}(0.25, 0.1)$ | 0.33[0.24, 0.61] |
| $\mu$ | $\mathcal{N}(0.15, 0.1)$ | 0.17[0.10, 0.23] |
| $\tau_l$ | $Exp(0.0001, 0.1)^{\dagger}$ | 0.7[0.005, 0.37] |
| $ur$ | $\mathcal{U}(0, 1)$ | 0.81[0.44, 0.99] |

The results point to a roughly 3 times higher transmissibility through sexual contact than via mosquito bites. We also see that about 20% of male cases go unreported.

## Discussion

We propose here a mathematical model for studying the sexual transmission of Zika. The model accommodates all known forms of Zika sexual transmission as well as vectorial transmission. Due to the current lack of knowledge about the relative effectiveness of the less common forms of sexual transmission, we have limited our analysis to the better understood transmission from men to women. This lack of a more complete knowledge of the ZIKV transmission cycle is the main limitation of our model. Once the actual transmissibilities of all modes of transmission are known, we will be able to benefit from the full potential of this model.

We have found that the sexual transmission of ZIKV can lead to a larger burden on women when compare to men (figure 1), simply due to the modified dynamics of transmission. Figure 5 shows the ratio *AR*_*W*_ (120)*/AR*_*M*_(120) in terms of the intensities of both sexual and vectorial transmission. This effect has been observed in Zika incidence data^10,13,22^ but it was frequently confounded with gender-related underreporting, but from our estimates of the male underreporting, we conclude that it can account for only a 20% drop, with the rest of the difference being explained by the excess in female cases due to sexual transmission.

This is true even when *ℛ*_0_ < 1, as illustrated in figure 5. Another interesting consequence of asymmetrical sexual transmission, i.e., it is easier for men to infect women than vice-versa, is that it allows us to differentiate its effects from those of a mere sexual bias in case reporting (It is know that women is more likely to report any illness than men), because the resulting dynamics are qualitatively different (figure 4). It may be difficult to differentiate these effects from noisy incidence data, but they are nevertheless different things and should be treated accordingly.

Figure 6 shows that for the reduced model, without homosexual and women to men transmission, vectorial transmission is necessary to allow crossing the epidemic threshold of *ℛ*_0_ = 1. It follows that if some positive flow of virus is possible directly from men to men or from women to men, epidemics will be possible without vectorial transmission, but although there are reported cases of both of these kinds of transmission^11^, 12, no observational evidence is available so far of sustained transmission without vectors. The fact that the correction factor *c* = 2.154 for male sexual contact heterogeneity is greater than 1, gives more importance to sexual transmission. Since men which have sex with men usually have substantially more sexual partners^23^, the amplification of the sexual transmission of Zika in this community can be quite large if this mode of transmission is included. Moreover, with effective male-to-male transmission, sustained transmission of ZIKV becomes possible without the vector.

**Figure 6.**
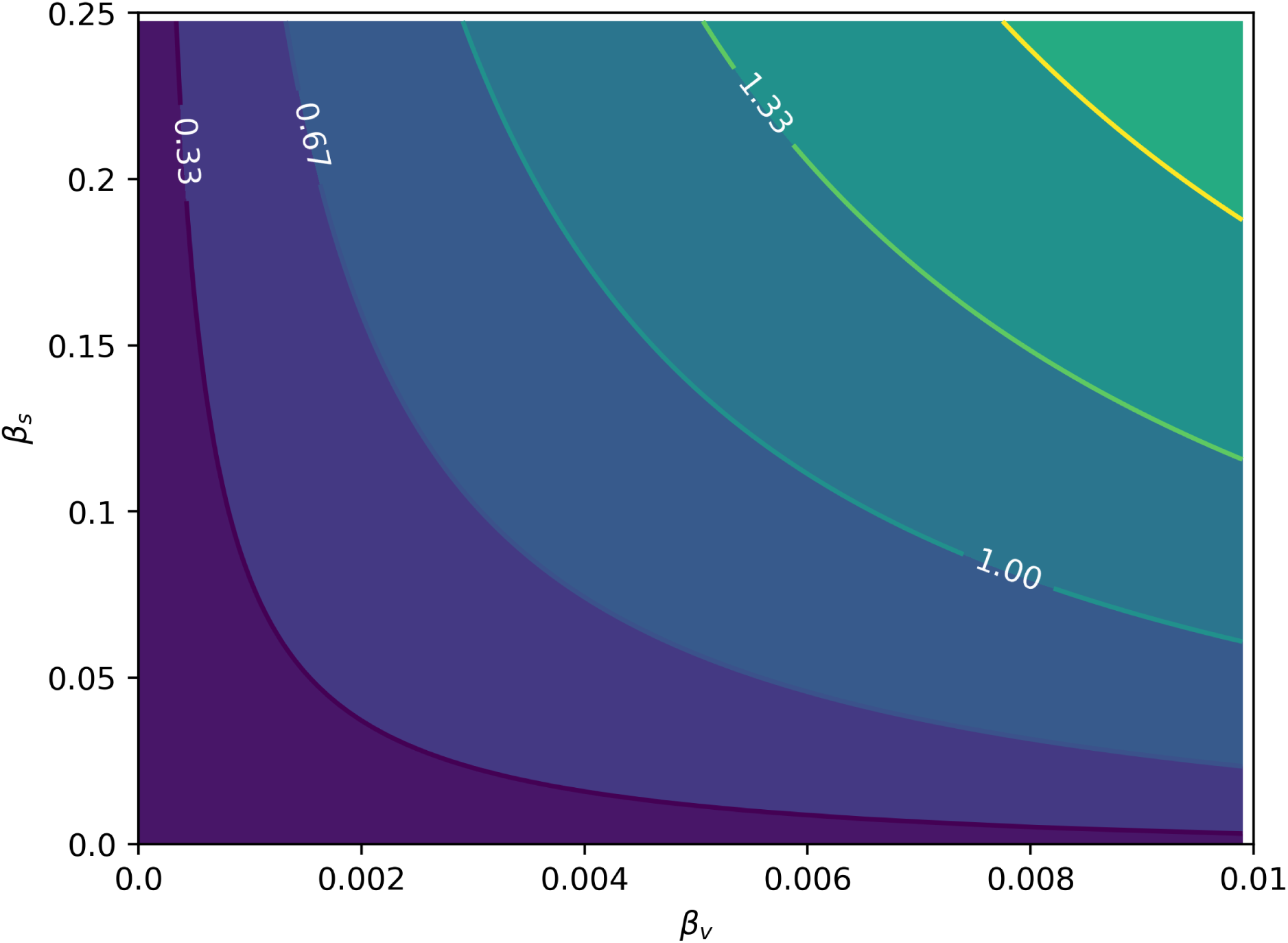
*ℛ*_0_ as a function of the relative intensities of sexual(*β*_*s*_) and vectorial(*β*_*v*_) transmissions. The *ℛ*_0_ values are already adjusted for the heterogeneity in sexual contact rates.

**Figure 7.**
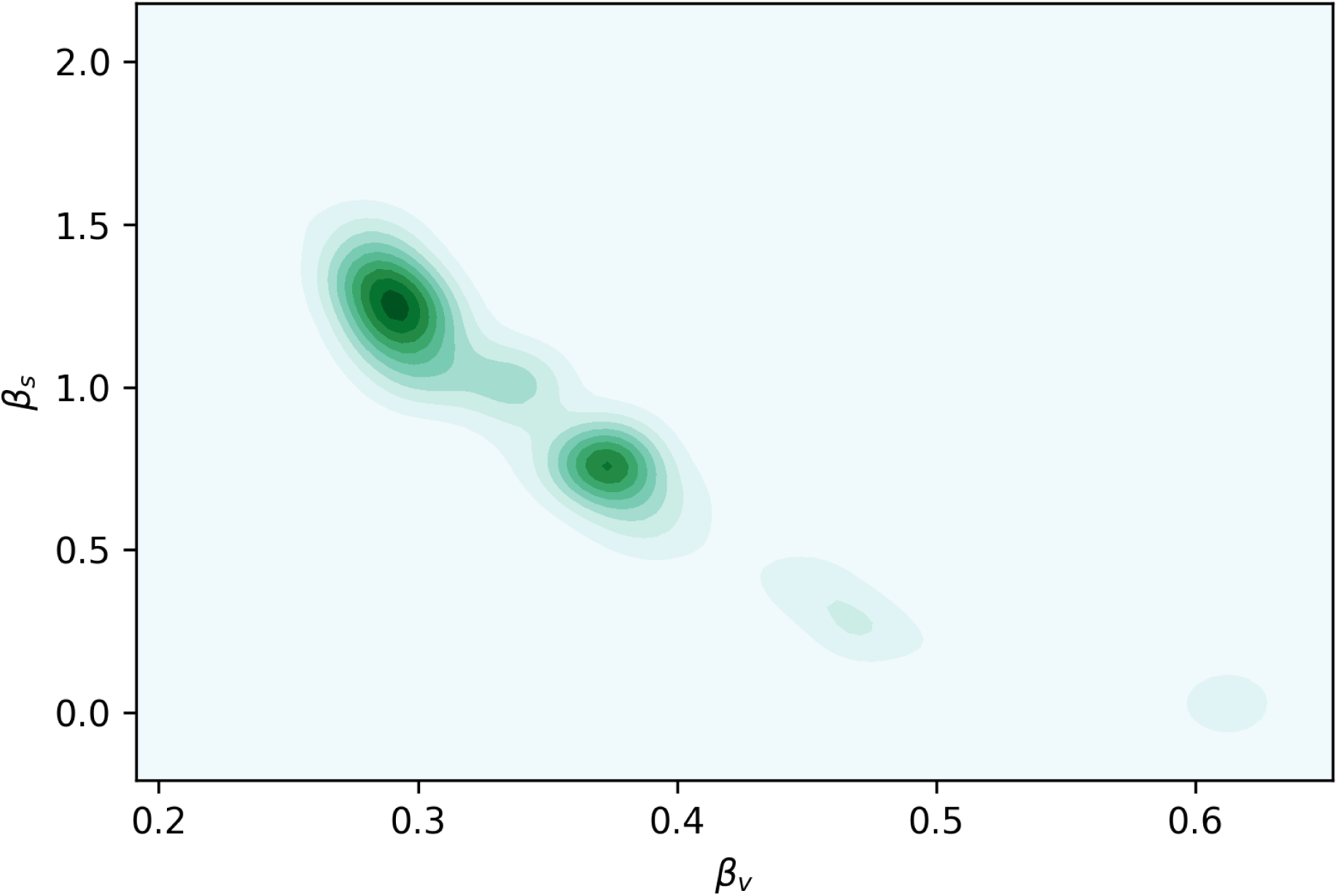
Joint posterior distribution of the transmission parameters *β*_*s*_ and *β*_*v*_.

**Figure 8.**
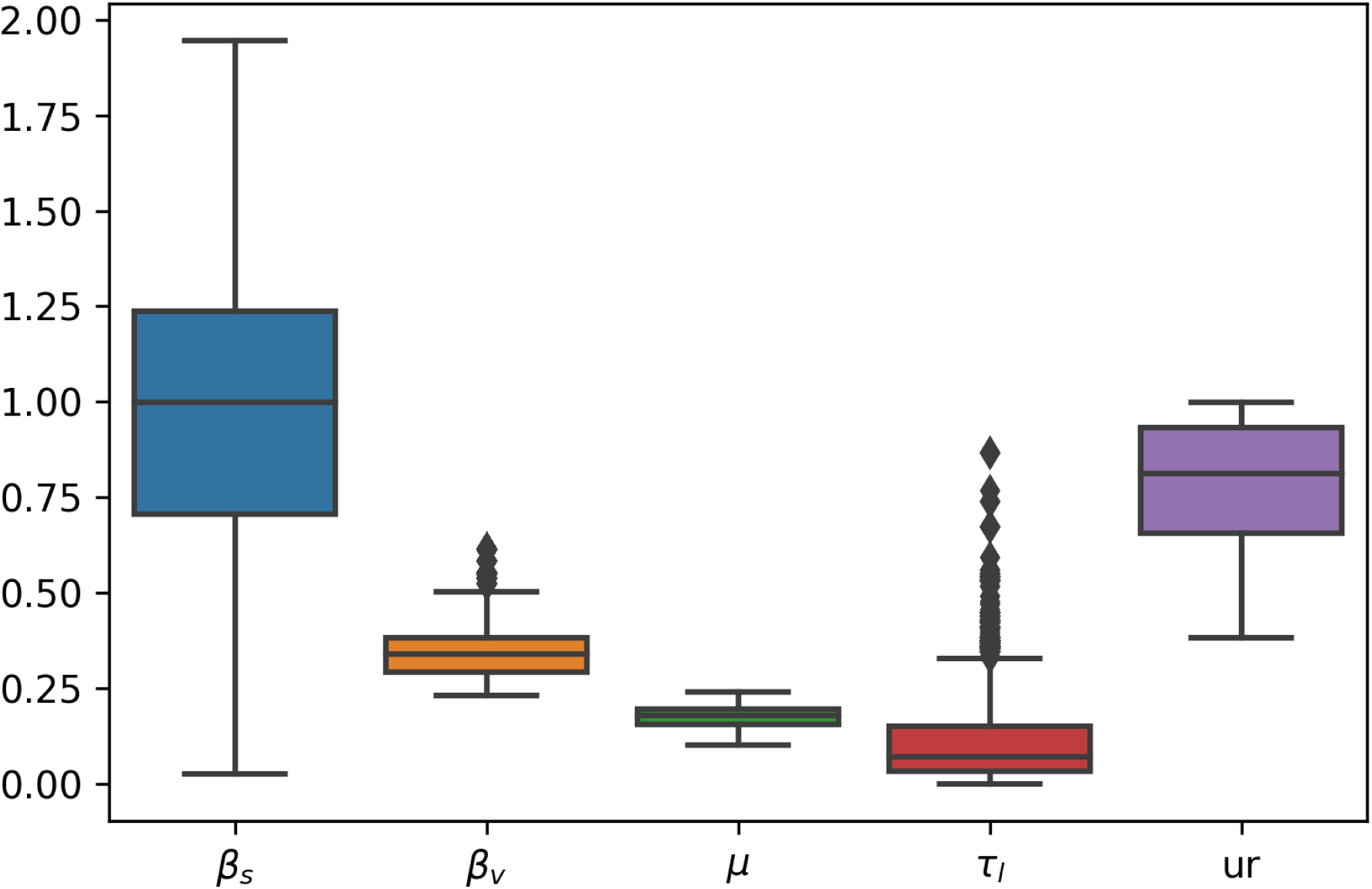
Boxplot of posterior distributions for all parameters estimated.

**Figure 9.**
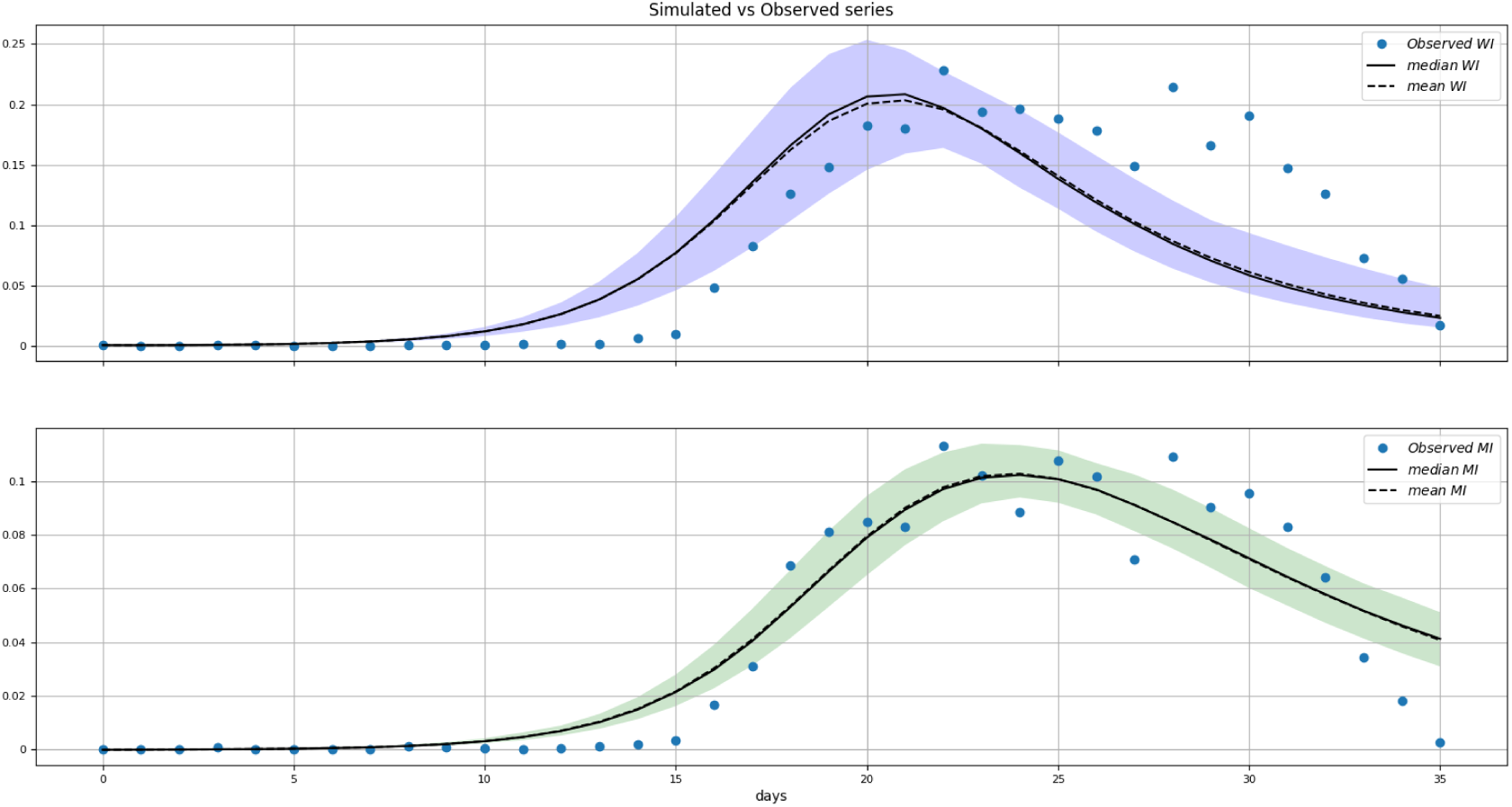
Posterior distributions of *W*_*I*_ (*t*) and *M*_*I*_ (*t*). blue dots are the data.

Another important factor in the determination of the long term dynamics of Zika in a population is the parameter *K*_*L*_ which represents how well men is capable of sexually transmitting ZIKV after the viremic period (figure 3), when compared to during it. It has been confirmed that some men can test positive for ZIKV RNA in the semen months after the viremic period^24,25^. Our model shows that until we can properly measure or estimate the sexual transmissibilities between humans in post-viremic stage, we must be prudent^26^ and recommend protected sex to men returning from endemic areas with or without symptoms.

## Acknowledgements (not compulsory)

The authors would like to thank Maria Soledad Aronna and Claudia Codeço for valuable comments on the manuscript.

## Author contributions statement

F.C.C., A.C.W.G.B. and K.G.S. conceived the Model, A.C.W.G.B. and K.G.S. implemented the model and calculated the *R*_0_ derived for the model, we decided investigate numerically the importance of the sexual transmission depending on, F.C.C., A.C.W.G.B. and K.G.S. analysed the model and numerical simulations. All authors reviewed the manuscript.

